# SweGen: A whole-genome map of genetic variability in a cross-section of the Swedish population

**DOI:** 10.1101/081505

**Authors:** Adam Ameur, Johan Dahlberg, Pall Olason, Francesco Vezzi, Robert Karlsson, Pär Lundin, Huiwen Che, Jessada Thutkawkorapin, Andreas Kusalananda Kähäri, Mats Dahlberg, Johan Viklund, Jonas Hagberg, Niclas Jareborg, Inger Jonasson, Åsa Johansson, Sverker Lundin, Daniel Nilsson, Björn Nystedt, Patrik Magnusson, Ulf Gyllensten

## Abstract

Here we describe the SweGen dataset, a high-quality map of genetic variation in the Swedish population. This data represents a basic resource for clinical genetics laboratories as well as for sequencing-based association studies, by providing information on the frequencies of genetic variants in a cohort that is well matched to national patient cohorts. To select samples for this study, we first examined the genetic structure of the Swedish population using high-density SNP-array data from a nation-wide population based cohort of over 10,000 individuals. From this sample collection, 1,000 individuals, reflecting a cross-section of the population and capturing the main genetic structure, were selected for whole genome sequencing (WGS). Analysis pipelines were developed for automated alignment, variant calling and quality control of the sequencing data. This resulted in a whole-genome map of aggregated variant frequencies in the Swedish population that we hereby release to the scientific community.

## Introduction

Whole-human genome sequencing (WGS) is now being performed at an unprecedented scale, due to technology developments making WGS affordable. The majority of this sequencing involves patient samples with a specific phenotype of possible genetic nature. Access to population-based reference control datasets, based on high-quality whole genome sequences, is important for identification of candidate disease causing genetic variants in clinical sequencing, for improved prediction of genetic effects in research studies, as well as for better imputation of samples typed on SNP arrays. International efforts to determine the genetic variability pattern in population-based samples, such as the 1000 Genomes project [1], has contributed important information, but due to the small sample size for many populations this provides merely an overview of the global pattern of variability. European populations differ in their genetic structure, and there is a need to assess the genetic variability at a more detailed level in individual populations. Such efforts have been initiated for example in The Netherlands, Denmark and Iceland [2–4]. Large-scale sequencing is also being performed in the UK but focusing on patient samples [5]. Similar WGS projects have also been initiated in many other parts of the world [6–9].

The genetic structure of the Swedish population was recently studied using genome-wide SNP data from 5,174 Swedes with extensive geographical coverage [10]. The results showed that there are pronounced differences, in particular between the northern and the remaining counties. The number of homozygous segments also showed large differences between southern and northern Sweden as well as between southern regions. Population sequencing efforts in Sweden have shown that a large number of the genetic variants in human populations have not yet been identified. For example, in sequencing 200 kb from 500 individuals from each of five local European populations, including northern Sweden, we have previously reported that 17% of single nucleotide variants and 61% of small insertion/deletions were not present in the public databases [11]. Most novel genetic variants showed a very limited geographic distribution, with 62% of the novel SNPs and 59% of novel insertion/deletion variants detected in only one of the local populations.

Thus, results from previous studies demonstrate that the genetic differentiation between local communities in Sweden is substantial when viewed on a European scale. The large number of population-specific low frequency SNPs and structural variants underscores the need to establish a database of genetic variants using whole human genome sequences for the Swedish population. It is well-known from chip array based genome wide association studies (GWAS) that genetic stratification within a population may introduce bias, particularly for rare variants. Careful handling of influences from genetic ancestry is therefore necessary to avoid false positives. The problem is likely to be more severe for sequence-based studies due to the multitude of low frequency variants. Comparisons between unmatched cases and controls will inflate levels of false positive and false negative findings [12]. Therefore, in order to make full use of WGS in association studies or in clinical diagnostics, there is a strong need to establish sufficiently large population-based databases for local populations. Finally, studies of human evolution and migrations have been brought to another level of understanding using WGS of both extant and archaic humans [13–15]. For these types of studies, population-based cross-sectional sampling provides the optimal strategy for estimating unbiased frequencies of individual variants as well as haplotypes.

Through the SweGen project described here, we make available a population-based high-quality genetic variant dataset for the Swedish population. The aims of this project were to identify a control cohort that reflects the genetic structure of the Swedish population, to employ WGS on these samples using Illumina X Ten sequencing, and to construct robust analysis pipelines that are capable of handling large-scale WGS projects. The genetic variant dataset based on these 1,000 individuals will enable scientifically sound whole genome sequence-based association studies for national patient cohorts studies, by providing data on well-matched national controls selected on the basis of the genetic structure of the population. It will also be an important resource in studies of familial heritable disorders and clinical sequencing, in particular for the filtering of genetic variants that are unlikely to have a pathogenic effect.

## Results

### Strategy for selecting individuals

The first aim was to identify a set of 1,000 individuals to be used for the construction of the SweGen reference cohort. Since this dataset will be used for a wide range of research projects as well as for routine clinical investigations, we decided to construct a population-based cross-sectional cohort that reflects the genetic structure of the Swedish population. As a first step we performed an inventory of available cohorts that fulfilled three criteria, i) population-based cross sectional cohorts (e.g. not focusing on specific outcomes), ii) SNP array data available to be able to study the genetic structure and, iii) DNA or blood samples available. A number of alternatives were identified and we decided to use samples from two of these; the Swedish Twin Registry (STR) [16] and the Northern Sweden Population Health Study (NSPHS)[17]. Both of these are population-based collections. STR is a national registry of Swedish born twins and the vast majority of samples (n=942) were taken from this resource. NSPHS was used to contribute a limited number of samples (n=58) from the very northern part of the country.

From the STR study, 10,000 samples had previously been genotyped using high-density SNP arrays and from NSPHS 1,033 individuals were available with similar data. Figure 1 shows a PCA plot with the first and second principal components (PCs) of these two Swedish cohorts, when viewed in the context of the diversity among the continental European populations included in the 1000 Genomes data [18]. The results indicate that the combined allele frequencies of known polymorphisms form a distinct pattern in the Swedish samples, thereby underscoring the need to generate a population-specific reference cohort. We identified a set of 1,000 samples for the SweGen dataset, 942 selected from STR and 58 selected from NSPHS and confirmed that they captured the diversity seen in the country. This cross-sectional selection of samples is likely to reflect the genetic variation that distinguishes the Swedish population from continental Europe. One caveat to this selection is that already established and well-characterized national sample collections such as STR and NSPHS, do not reflect recent migration patterns. Therefore, the SweGen control cohort is likely to reflect the genetic structure of Swedish individuals that have been present in Sweden for at least one generation.

**Figure 1.**
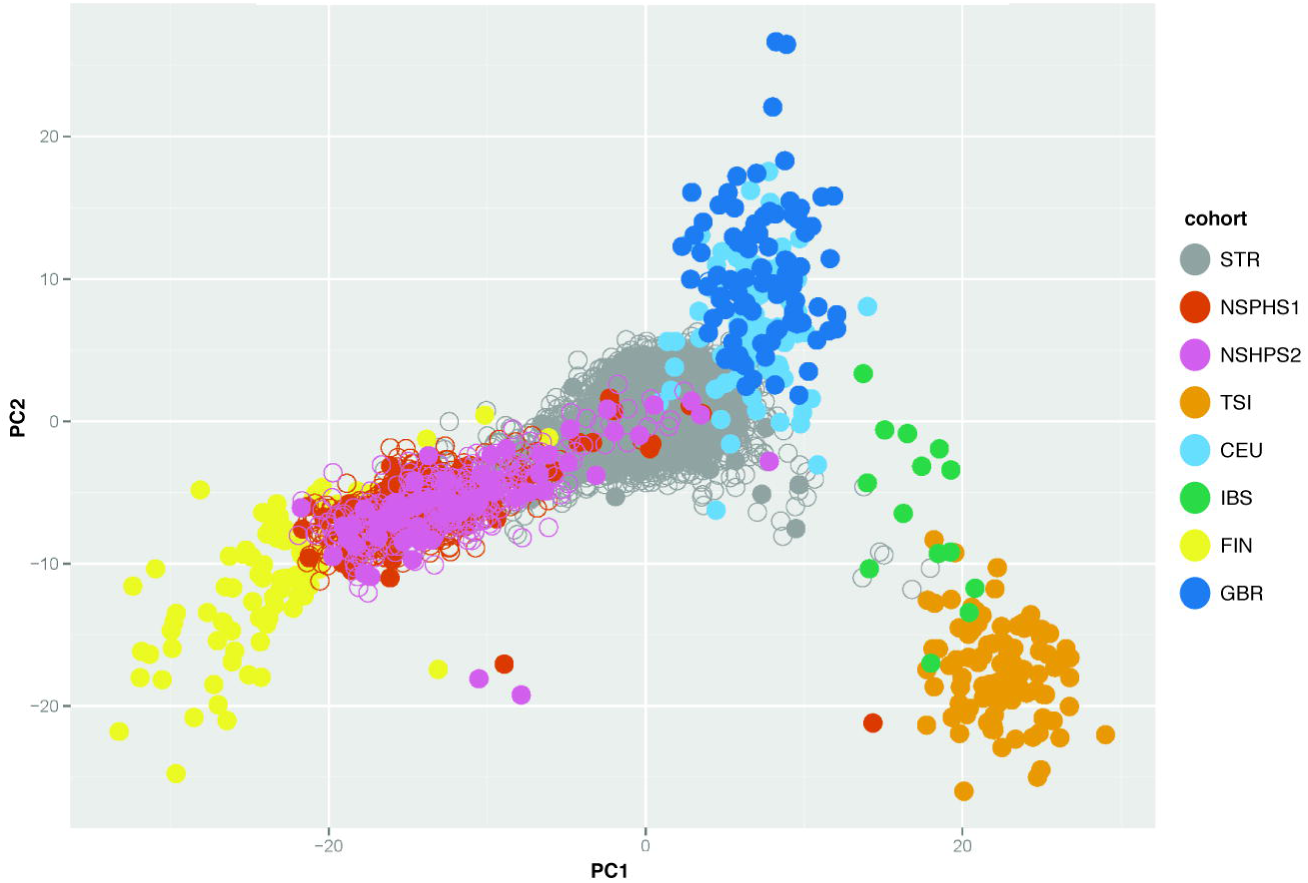
Genetic variation within Sweden related to European populations. The plot shows the two first principal components from a PCA where samples from the Swedish Twin Registry (STR) and the Northern Sweden Population Health Study (NSPHS1 and NSPHS2, collected in two different phases) are compared to available data from European 1000 Genomes populations (CEU: Utah Residents with Northern and Western Ancestry, FIN: Finnish in Finland, GBR: British in England and Scotland, IBS: Iberian Population in Spain, TSI: Toscani in Italia). The analysis is based on 19978 autosomal SNPs, selected to fulfill the following i) must be present (by name) on SNP chips Affymetrix 5.0, Affymetrix 6.0, Illumina OmniExpress and in the 1000 genomes phase 1 EUR dataset ii) per SNP missingness <= 2% in the combined sample iii) minor allele frequency >= 5% in the combined sample iv) pairwise LD (R^2) <= 0.05 with other SNPs within a 1000-SNP window and v) not in any of a list of regions known to harbor strong long-range LD.SNP weights were defined by running the PCA in European 1000 Genomes samples, and then projecting STR and NSPHS into the resulting space defined by the analysis.

### Results of pilot experiments

The 1,000 Swedish individuals were sequenced on the Illumina X Ten system at two different sites; NGI Uppsala (NGI-U) and NGI Stockholm (NGI-S) (see Methods). Prior to sequencing the entire dataset, two small pilot studies were conducted to validate the quality of the WGS data. In the first pilot experiment, performed at NGI-S, two independent library preparations from the same sample were analyzed. Sequencing and analysis of the two replicated libraries resulted in approximately 3.86M single nucleotide variant (SNV) positions, where the duplicate sample had a non-reference genotype called for either of the library preparations. Of these, 90,446, or 2.3%, had discordant genotypes, resulting in a genotyping concordance of 97.7%

In a second pilot study, 8 samples were sequenced in parallel at NGI-U and NGI-S to study potential biases between the two sites. Our results revealed that out of an average of 8,056,903 genotyped variant sites for each of the 8 sample pairs, 7,942,822 genotype calls on average overlap between the two replicate samples, prior to any filtering. The genotype concordance between the samples was 97.5% on average. Since very similar concordance results were obtained from the first pilot study, we conclude that the site-effects between NGI-U and NGI-S are negligible. After variant quality score recalibration (VQSR), the number of overlapping genotypes, passing filters, was on average 7,448,632 sites per sample pair (93.8% of the number of sites pre-VQSR) and the genotype concordance was 98.6%, on average. The genotype concordance of variants pre- and post-VQSR filtering shows the effectiveness of this approach, as more than half of the discordant sites (104,981/196,822 = 53.4%) are removed, while filtering only 6.2% of the total number of sites. To assess whether there was any systematic lack of coverage in coding regions for the 8 pilot samples, we listed all exons with less than 50% sequence coverage in more than half of the samples. A total of 31 exons from 8 gene loci were found to be less than 50% covered. Only one single exon had less than half of its positions covered in all 8 samples.

### Analysis and quality control of the WGS data

Of the 1000 samples, 509 were sequenced at NGI-S and 491 at NGI-U. For the management, alignment and SNV analysis of the WGS data, we developed a robust and efficient pipeline that was used for all samples (see Figure 2). Briefly, this pipeline aligned the reads to the human reference genome (GRCh37/hg19 assembly version), processed the alignments according the GATK best practice recommendations [19] and carried out variant calling using the GATK HaplotypeCaller [20]. This workflow was run using Piper [21].

**Figure 2.**
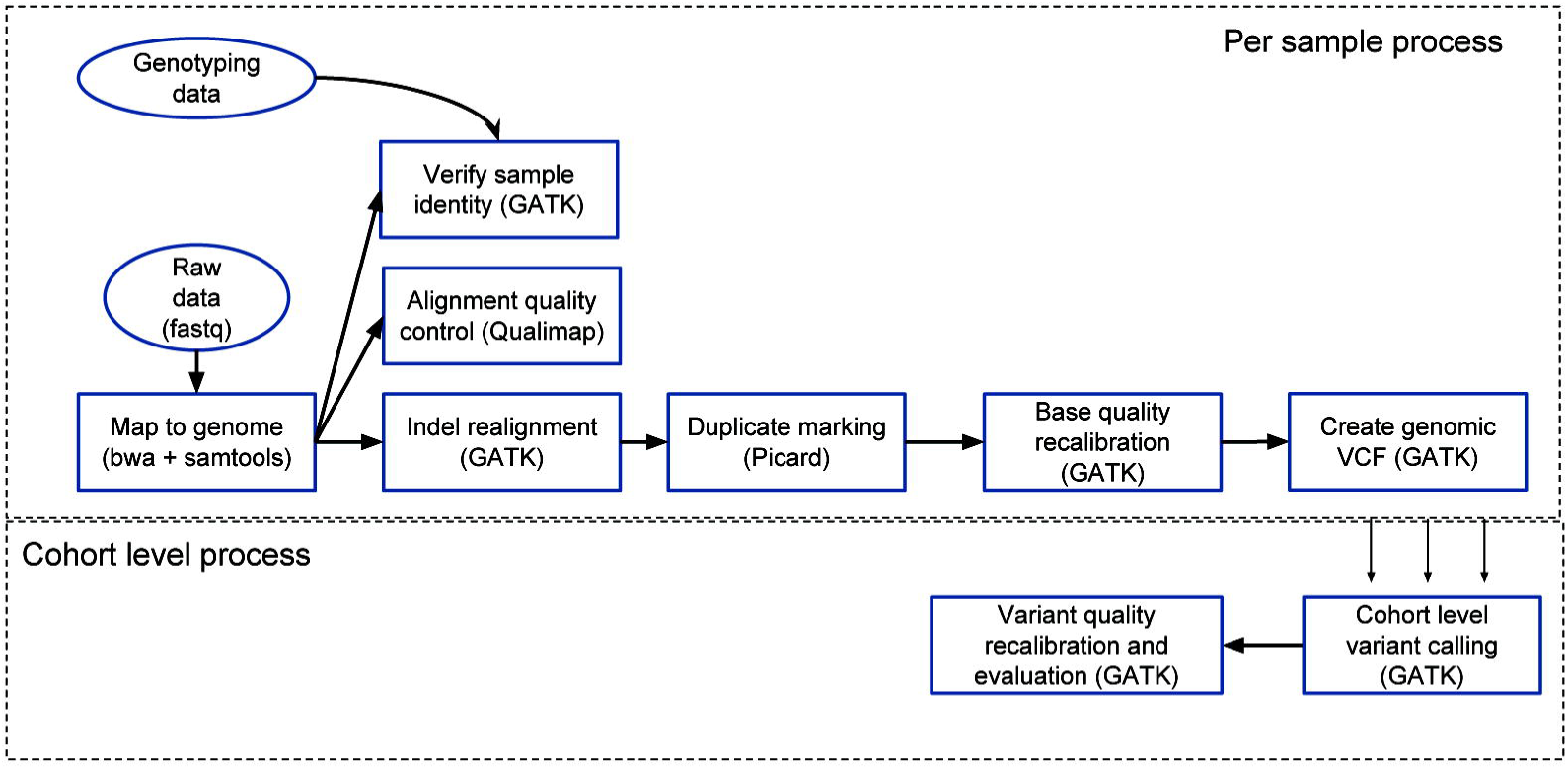
Overview of workflow for alignment and SNV detection. The process has two phases, first each sample is processed individually and then the entire cohort is processed together. The first phase begins by aligning the raw reads to the reference genome using bwa, converted the resulting alignments to bam format, and sorting and indexing them using samtools. Preliminary sample identity is verified by checking concordance with genotyping data and alignment quality is assessed using Qualimap. Once all alignments from a sample has been merged, they are processed according the GATK Best practice workflow, with indel realignment, duplicate marking and base quality score recalibration, before using the GATK Haplotypecaller to create genomic VCF files (GVCF). The second phase is carried out on a cohort level. The GVCF files are combined SNVs and INDELS are called. This is followed by variant quality recalibration and recalibration. Finally quality control metrics and population statistics are computed for the final call set.

A series of analyses were performed to assess the data quality. To ensure that no sample mix-ups occurred during the sequencing procedure, all samples were independently genotyped at a limited number of SNP positions and checked for concordance with the WGS data. We also performed a kinship analysis to ensure that no sample was represented in duplicate in the final dataset. Further, we performed a PCA on all 1,000 samples to ensure that there were no unexpected outliers or obvious biases between samples sequenced at the two different sites (Figure 3). After having performed the basic quality control steps, we generated a file containing the SNP frequencies observed in the 1,000-sample dataset. The resulting frequency file is available for the research community both as a flat text file and through a local installation of the ExAC [22] browser.

**Figure 3.**
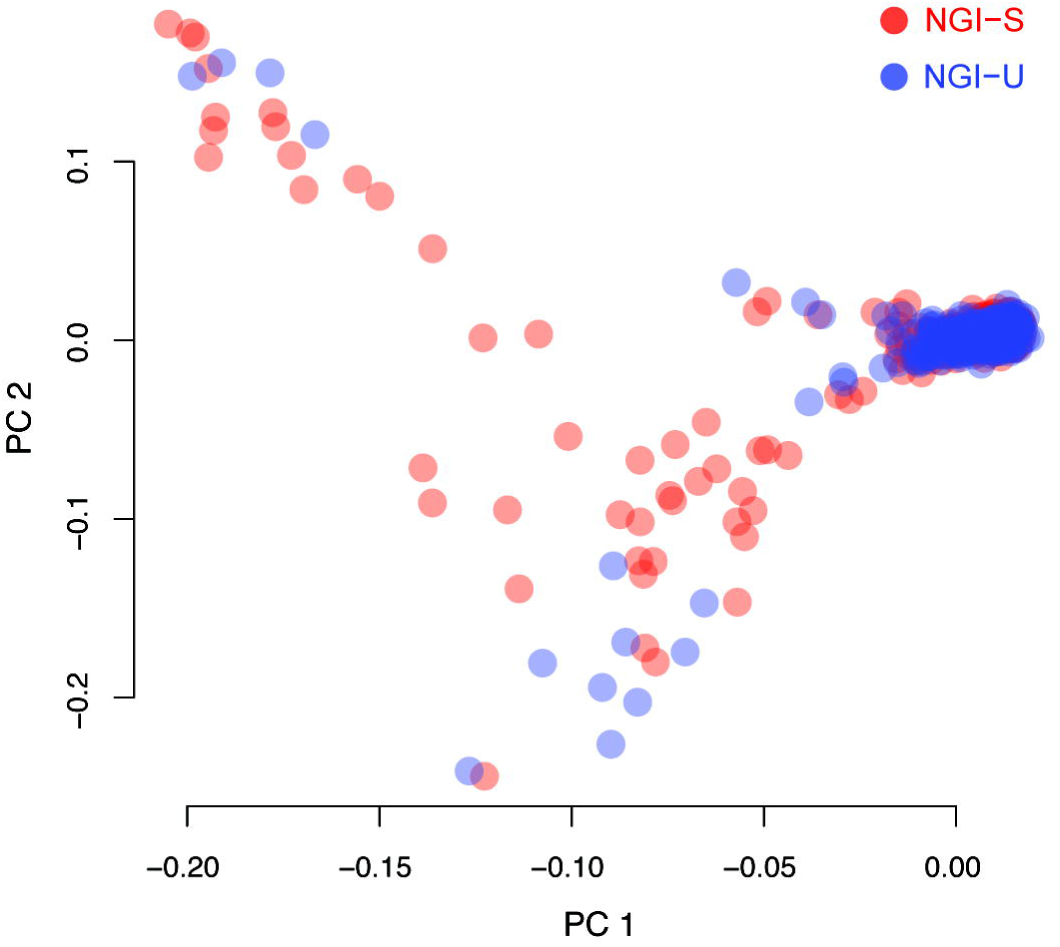
PCA of variation between sequencing sites in the 1000 WGS samples. 509 of the SweGen samples were sequenced at NGI-S (red) and 491 samples at NGI-U (blue). The PCA plot shows the distribution of genetic variation based on ~650k common SNPs from the Illumina OmniExpress chip. There are no clear biases detected in the first two principal components when comparing the two sites.

### Overview of the SweGen dataset

An overview of the WGS data for all 1,000 samples in the SweGen dataset is shown in Table 1. The average autosomal coverage after duplicate removal was 36.7X, with values ranging between 20.2X and 97.6X for individual genomes. A total of 29.2 million SNP sites and 3.8 million indel sites were detected in the 1000 samples. Of these, 8.9 million SNPs and 1.0 million indels were not present in version 147 of dbSNP [23], a database that includes information from phase 3 of the 1000 Genomes Project and many other sources. 7,198,518 of the novel variants were detected in only one of the samples, thereby resulting in an average of 7,199 novel variants per individual. As a comparison, a recent publication on 10,000 whole genomes detected 8,579 novel variants per individual [6]. For the novel genetic variants, 23,396 SNPs and 3,239 indels were found to have a direct effect on the amino acid sequence of some protein (i.e. non-synonymous, stop-gain, stop-loss, frame shift or splicing). Our results thus indicate the presence of a substantial number of genetic variants in the Swedish population that are not represented in current databases, some of which could have a direct effect on a protein.

**Table 1.** Overview of the WGS data for the 1000 Swedish samples

|  |  |
| --- | --- |
|  | SweGen dataset<br>(STR+NSPHS) |
| Nr samples: | 1000 |
| Avg coverage:<br>[min /max] | 36.7<br>[20.2 / 97.5] |
| Total nr SNP sites:<br>[not in dbSNP147] | 29,162,141<br>[8,856,354] |
| Total nr indel sites:<br>[not in dbSNP147] | 3,825,043<br>[1,001,047] |
| Ts/tv ratio: | 2.01 |
| Avg homoz SNPs:<br>[min/max] | 1,486,648<br>[1,332,739 / 1,578,505] |
| Avg heteroz SNPs:<br>[min/max] | 2,366,095<br>[2,190,824 / 2,603,755] |
| Avg singleton SNPs:<br>[min/max] | 10,975<br>[1,030 / 31,087] |

## Discussion

We have generated a map of the landscape of genetic variability in the Swedish population. The dataset is constructed from a population-based selection of individuals, taking into account the main aspects of the genetic structure of the Swedish population. The individuals represent a cross-section of the population; meaning that it contains the general population prevalence of diseases, and no metadata has been used in the selection of individuals to be included. Also, we do not have information on disease outcomes for the individuals included. This implies caution that the dataset likely includes mutations that are associated with, or causative of, disease.

A limitation of the SweGen dataset is that it lacks representation of the genetic constitution of individuals arriving recently to Sweden. About 10% of the present Swedish population has an origin in a country outside Europe, and have only recently settled in Sweden. This fraction has been further affected by the refugee situation in the last year, many of which are seeking permanent residence in Sweden. In order to appropriately include also the genetic diversity of these groups, specific additions to the control cohort may be needed, although it might be more feasible to rely on similar genome initiatives carried out in the different regions of origin, since they may be more appropriate designed, depending on the ethic diversity in each country. In spite of these limitations the database represent a valuable annotation of the genetic variability pattern in the majority of the Swedish population.

In this first release, the data is presented as a joint frequency table, annotating the frequency of SNPs and small insertion/deletion variants at each position across the genome. This is aimed at users in need of identifying and filtering out non-pathogenic population-specific low frequency variants from their patient sequences. However, this type of data can also be used for WGS-GWAS, although these analyses are more powerful when based on individual genotype information. The individual genotype information will be made available at a later stage to individual projects granted permission to access the data. This will enable for more powerful studies the underlying genetic structure of the population, including haplotype-based analyses and imputation.

The size of the control cohort is a balance between the statistical power to identify genetic variants and the cost for WGS. The control database of 1,000 individuals has sufficient size to include genetic variants down to a frequency of around 1 %, although only a subset of all variants are present at that frequency in the population. In terms of the power in association studies, sample sizes on the order of 10^3^ are needed to detect effects of low frequency variants [24]. Given that the cost of WGS continues to plummet, the data set could be extended to increase our ability to detect a higher fraction of the low frequency genetic variants.

Our results indicate that the SweGen dataset contains genetic variation that is not represented in the European populations included in the 1000 Genomes Project. By contributing information on this previously unreported genetic variation, as well as a building a reference dataset for the entire population, we believe that the SweGen dataset will provide a valuable national resource for genetics research and clinical diagnostics.

## Methods

### DNA samples and ethical consent

The Swedish Registry (STR) was established in the 1960s and, at present, holds information on 85 000 twin pairs, both monozygotic and dizygotic. DNA from blood or saliva have been collected and extracted from 55,000 of these individuals. 10,000 participants (one per twin pair for monozygous twins) in the TwinGene project were subjected to high-density SNP array typing. TwinGene is a nation-wide and population based study of Swedish born twins agreeing to participate. The TwinGene sample collection represents the Swedish geographic population density distribution. In our initial PCA all 10,000 individuals were included, and from this analysis, 942 unrelated individuals were selected for WGS, mirroring the density distribution. The Twingene study was approved by the regional ethics committee (Regionala Etikprövningsnämnden, Stockholm, dnr 2007-644-31, dnr 2014/521-32).

The Northern Sweden Population Health Study (NSPHS) is a health survey in the county of Norrbotten, Sweden to study the medical consequences of lifestyle and genetics. Blood samples were collected (serum and plasma) and stored at −70°C on site. DNA was extracted using organic-extraction. 58 samples were selected based on PCA. The NSPHS study was approved by the local ethics committee at the University of Uppsala (Regionala Etikprövningsnämnden, Uppsala, 2005:325 and 2016-03-09). All participants gave their written informed consent to the study including the examination of environmental and genetic causes of disease in compliance with the Declaration of Helsinki.

### Library preparation and Illumina sequencing

Library preparation and sequencing was performed by the National Genomics Infrastructure (NGI) platform in Sweden, at the Genomics Production site in Stockholm (NGI-S) and SNP&SEQ facility in Uppsala (NGI-U). The 1000 samples were divided between NGI-S (509 samples) and NGI-U (491 samples) and library preparation and sequencing were performed independently by each facility. Sequencing libraries were fragmented with Covaris E220 to 350 bp insert sizes and were prepared from 1.1µg/1µg DNA (NGI-S and NGI-U, respectively) using the TruSeq PCRfree DNA sample preparation kit according to the manufacturer’s instructions (guide#15036187). The protocols were automated using an Agilent NGS workstation (Agilent, CA, USA) and a Biomek FXp (Beckman Coulter Inc.) at NGI-S and NGI-U, respectively. Clustering was done using cBot. Paired-end sequencing with 150bp read length was performed on Illumina HiSeqX (HiSeq Control Software 3.3.39/RTA 2.7.1) with v2.5 sequencing chemistry. A sequencing library for the phage PhiX was included as 1% spike-in in the sequencing run.

### Alignment and variant calling

Raw reads were aligned to the 1000 Genomes (b37) fasta reference, using bwa mem 0.7.12 [25]. The resulting alignments were sorted and indexed using samtools 0.1.19 [26]. Alignment quality control statistics were gathered using qualimap v2.2 [27]. Alignments for the same sample from different flowcells and lanes were merged using Picard MergeSamFiles 1.120 (broadinstitute.github.io/picard). The raw alignments were then processed following the GATK best practice [19] with version 3.3 of the GATK software suite. Alignments were realigned around indels using GATK RealignerTargetCreator and IndelRealigner, duplicate marked using Picard MarkDuplicates 1.120, and base quality scores were recalibrated using GATK BaseRecalibrator. Finally gVCF files were created for each sample using the GATK HaplotypeCaller 3.3. Reference files from the GATK 2.8 resource bundle were used throughout. All these steps were coordinated using Piper v1.4.0 [21].

### Population calling and generation of variant frequency file

Population calling (i.e., joint genotyping) was executed on the whole cohort of 1000 samples as recommended by GATK guidelines. This produced a single VCF file containing all variants identified in the 1000 sequenced individuals. Due to the large number of samples, 5 batches of 200 samples each have been merged into 5 separate gvcf files using GATK CombineGVCFs. The 5 individual gvcf files were used as input for GATK GenotypeGVCF. Subsequently, SNPs and Indels were extracted from the resulting gVCF files and GATK VQSR steps executed on both sets of variants (i.e., GATK VariantRecalibrator and GATK ApplyRecalibration). In order to reduce the overall time the first two steps (GATK CombineGVCFs and GATK GenotypeGVCF) were run on 60Mbp non-overlapping segments of the Human Genome.

## Data availability

The SweGen variant frequency data is available from the site https://swefreq.nbis.se. Individual positions can be browsed using a local installation of the ExAC browser [22]. A flat file containing frequency data for the whole genome is available for academic use upon registration and agreement to terms and conditions for data download.

## Acknowledgements

This work was funded by Science for Life Laboratory (SciLifeLab) as a National Project, supported by the Knut and Alice Wallenberg Foundation (2014.0272), and The National Research Council (PI:UG). Illumina sequencing was performed by the National Genomics Infrastructure (NGI), which is hosted by SciLifeLab in Stockholm and Uppsala. Computational resources from UPPNEX were used for data analysis.

